# Community ecology dynamics on reefs in Fiji are dependent on soft coral density and depth

**DOI:** 10.1101/2025.08.15.670538

**Authors:** Nile P. Stephenson, Katie M. Delahooke, Charlotte G. Kenchington, Tasnuva Ming Khan, Lea Katz, Jone Waitaiti, Alice A. Ball, Victor E. Bonito, Emily G. Mitchell, Andrea Manica

**Affiliations:** Department of Zoology, University of Cambridge, UK; University Museum of Zoology, Cambridge, UK; Department of Earth Sciences, University of Cambridge, UK; British Antarctic Survey, Cambridge, UK; Marine Biology Lab, Université Libre de Bruxelles, Brussels, Belgium; Viani Bay Dive School, Vanua Levu, Fiji; Reef Explorer Fiji, Votua Village, Fiji

**Keywords:** Soft coral, Spatial ecology, Bayesian network inference, Fiji, Mutualism

## Abstract

Some soft corals (order Alcyonacea) have the potential to increase in prevalence on tropical coral reefs as the severity of anthropogenic climate change increases. While soft corals are therefore an increasingly important component of reef ecosystems, little is known about their ecological role on coral reefs and their influence on community dynamics and diversity. We used Bayesian Network Inference to identify the relationships among benthic taxa across sites with varying degrees of soft coral dominance on the Great White Wall, Fiji, and then employed spatial point process analysis to reveal the ecological processes behind these associations. We found that depth was the dominant driver of community dynamics, and that white Nephtheidae soft corals were negatively associated with scleractinian corals and positively associated with algae due to a facilitative mutualism – possibly due to soft corals reducing grazing pressure. Our results show distinctions in reef benthos and ecological dynamics between scleractinian- and soft coral-dominated reefs. We found that diversity levels were significantly lower on soft coral than scleractinian reefs, potentially highlighting the risk of a loss in benthic diversity on reefs where soft corals replace scleractinians.

## Introduction

Tropical coral reefs are among the most biodiverse ecosystems on Earth (1) but are under threat from anthropogenic climate change (2–5). As the frequency and severity of threats such as marine heatwaves, ocean acidification, pollution, and overharvesting increases, coral reef communities are expected to change in composition (5–7), with an increase of organisms that are more resistant to the effects of climate change (8–11). Coral reefs are dominated by scleractinian corals, but soft corals (order: Alcyonacea) are a common occurrence (12,13) and sometimes form carpets of local dominance (14,15). While some of the soft corals are susceptible to the same pressures as scleractinian corals (11,16), many are resistant to, or benefiting from, changing climate conditions because of physiological adaptations (e.g., nitrate eutrophication in *Xenia*, tolerance of acidification in *Sarcophyton* and *Veretillum*) or warming-resistant symbiont-host dynamics (6,7,11,17,18). As such, it is possible that soft corals play a major role in coral reefs under future climate scenarios (6,19,20).

The biodiversity benefits of scleractinian coral reefs are emergent properties of both the composition and structure of reefs (21,22), and their ecological dynamics (23). It is unclear how ecological dynamics will change if soft corals become more prevalent (19) because much of the biodiversity generated by scleractinians is attributable to ecosystem engineering (22,24). Many soft corals do not produce such benefits because they provide only temporary habitat due to periodic retraction of their bodies and only over their own lifetimes (25). It is therefore important to understand the ecological dynamics of soft coral reefs to understand how reefs may change as soft corals increase in prevalence, and how these changes will impact reef biodiversity.

Ecological interactions within communities can be identified via Bayesian Network Inference (BNI), which allows the conditional dependencies between variables (in an ecological context, species or environmental characteristics) (26–29). In BNI, nodes (species, environment) are connected by edges (ecological associations). Edges have a sign (positive, negative) denoting the nature of the interaction and the influence score denotes the interaction’s strength (30). BNs are scale-agnostic, so variables can include biotic (species, genes) and abiotic (space, time, depth) variables, and biotic variables at different taxonomic levels (27,28,30,31).

While BNs can resolve the sign and strength of associations between variables, their ecological drivers cannot be determined by BNs alone (29,31). Pairwise ecological processes that drive associations within sessile species can be inferred using spatial point process analysis (SPPA) because their spatial patterns reflect the underlying processes that have occurred throughout an organism’s life (32–34). Points (organism locations) of a pair of species can be: i) independent, reflecting no associations; ii) aggregated, reflecting aggregation of points between populations (denoting e.g., mutualisms) or; iii) segregated, reflecting hyperdispersion of points of each population (denoting e.g., mortality incurred by competition) (34,35). Each of these patterns (independence, aggregation, segregation) can occur across multiple spatial scales and at different strengths. For instance, segregation at small scales and aggregation at large scales may indicate the influence of resource competition at small scales and shared habitat preferences at larger scales (35). The most likely underlying processes can be assessed by model-fitting of known processes and thus inference of those underlying processes (34,36).

In this study, to assess how ecological dynamics vary with soft coral dominance, we applied BNI and SPPA sequentially to three sites on the Great White Wall (GWW) reef, Fiji. BNI was performed on all sites together to elucidate the core ecological associations on the GWW, and subsequent SPPA was used to interrogate the dynamics underpinning these associations. We used linear models to understand how changing soft coral dominance influenced biodiversity.

## Materials and methods

### Field site

In this study, we focussed on the GWW, a vertical section of barrier reef in the Somosomo Strait, Fiji. The main GWW (mGWW) is an ideal site to study soft coral ecological dynamics because it has a large spatial extent dominated by a carpet of the white Nephtheidae (37). To understand how ecological dynamics differed between reefs of variable soft coral dominance, we added two more sites: the adjacent wall reef (AWR), also a wall reef but with fewer white Nephtheidae, more scleractinians, and a mixed filter feeding community; and the barrier reef flat (BRF), a shallow reef dominated by scleractinians and colourful *Dendronephthya* soft corals.

Across the three sites, we identified seven abundant benthic morphospecies: i) white Nephtheidae soft corals, ii) colonial hydroids (possibly *Myriopathes*), lemon sponges (*Leucetta chagosensis*), colourful *Dendronephthya* soft corals, *Halimeda* algae, adult *Tubastraea* sp. (an azooxanthellate stony hexacoral), a “hard coral” functional group which included all zooxanthellate, reef-building corals (e.g., *Acropora*, *Pocillopora*, *Porites*, *Favites*), and an encrusting sponge group, including sponges that encrusted atop substrate (e.g., *Monanchora, Aplysinella, Callyspongia*). Examples shown in Fig. S1d-k. Further benthic taxa found within the study area but not in sufficient abundance (n<100) for the analyses used here included: i) sponges: *Stylissa, Rhabdastrella, Neopetrosia*, and the endolithic sponge *Siphonodictyon*; ii) soft corals: *Sarcophyton*, *Cladiella*, *Klyxum*, and *Chironephthya*; iii) whip gorgonians; iv) sea fans, possibly *Annela, Melithaea, Subergorgia*, and *Acanthogorgia*); v) Stylasterids; vi) tunicates (*Polyandrocarpa*, *Didemnum*); vii) holothurians (*Pearsonothuria*); viii) crinoids (*Anneissia*); ix) molluscs including mussels (*Pedum, Tridacna*), sea slugs (*Phyllidia*), and nudibranchs (*Chromodoris lochi, Pteraeolidia semperi*). In this study, we identified organisms to a morphospecies level due to the high degree of cryptic diversity present in soft corals, and because collection of biological materials for finer resolution identification was not possible.

### Data collection

We used structure-from-motion photogrammetry to construct maps of each GWW site following (37). Video footage was collected in a lawnmower pattern and then split into frames at one frame/second using FFmpeg (38), and blurry frames removed. For each site, frames were aligned to form 3D models in Agisoft Metashape v2.1.1 and laser dots as a scale. A 2D “orthomosaic” photo map was generated from each 3D model and organisms were outlined and identified from each map with the morphospecies and location recorded. We used additional video and photographs to check morphospecies identification. The marked-up maps were extracted using a custom script (available at https://github.com/nis38/NPS_dex.git, adapted from (39)) in R v4.2.2 (40).

### Bayesian network inference

We performed BNI in software banjo (Bayesian network inference with Java objects) v2.2.0 (for details see (28,41); Supplementary Methods). We subset our spatial maps for BNI into 2m^2^ quadrats *cf.* (29), and converted counts into density to account for to the uneven edges of the mapped areas. Each quadrat was assigned a depth corresponding to either the BRF (12m) or the depth bins identified in (37): shallow (17 – 23.5m), intermediate (23.5 – 28m), and deep (28 – 33.5m). Depth bin definitions were dictated by variation in GWW topography; the intermediate depth bin contains four shallow caves (Fig. 2 in (37)), which are areas that interact with fluid flow (and thus food and light availability), whereas these caves were absent from shallow and deep bins.

### Spatial point process analysis

To understand underlying processes that result in the BNI edges, we performed bivariate SPPA for taxa with sufficient sample sizes (n>100 each). The pair correlation function (PCF) was used to test model fits between the observed point pattern of each edge against known spatial patterns (36). PCFs represent how the density of points (e.g., corals) at a given distance *r* from a given point change as a function of distance from that point (36). While inferring ecological processes from patterns is difficult because of the possible superposition of multiple processes (i.e., the overlaying of intra- and interspecific interactions or variable environmental preferences) and equifinality (where different processes generate similar spatial patterns) (34,45–47), the application of complimentary statistical techniques can disentangle processes underpinning different ecological patterns, particularly across differing spatial scales (34,46). We performed spatial analysis in Programita (v Novembre 2018) (48) (see Supplementary Methods).

### Rare morphospecies and diversity analysis

In most ecosystems, the majority of species are rare (57,58) and the identity of rare species may vary with depth or the dominant morphospecies (e.g., scleractinian *vs.* soft corals). For the BNI and SPPA, rare species could not be included in our analysis (28) so, to investigate the possible effects of rare taxa, we removed all common morphospecies for each quadrat to create a dataset of rare morphospecies. We used a linear discriminants analysis (LDA) with depth as a predictor variable and density of each rare morphospecies as response variables. An LDA is a supervised ordination method whereby clustering is dictated by the predictor variable (59). The accuracy of the LDA was tested using a training-testing split of 60:40, whereby 60% of the data was used to construct the LDA, and the results of the LDA were used to predict the remaining 40% of the data.

In order to assess the impacts of the soft corals and depth on the diversity of each site, Simpson’s index (60), Shannon evenness (61,62), and Fisher’s alpha parameter (63) were calculated for each quadrat. Each metric was used as a response variable in separate linear mixed models using the R package lme4 (64) with depth bin and density of white Nephtheidae as fixed response variables and site (BRF, GWW, and AWR) as a random variable in each model.

## Results

White Nephtheidae were the dominant morphospecies on the intermediate and deep depth bins of mGWW (87% relative abundance) and in the intermediate depth bin of the ARW (80% relative abundance). Mixed scleractinian and soft corals were present in the shallow mGWW (6% scleractinian, 66% soft coral) and the shallow AWR (30% scleractinian, 9% soft coral). The BRF was evenly split between scleractinians (47%) and soft corals (48%, mostly *Dendronephthya*) (Table S1).

### Bayesian network inference

Nine nodes (eight morphospecies and depth) were included in the best BN with a linkage density of 0.78. Depth was the dominant node directly influencing the density of the white Nephtheidae and hydroids positively, and *Dendronephthya* negatively. Depth indirectly influenced all other nodes apart from lemon sponges, which had no edges. The white Nephtheidae positively influenced *Halimedia* algae and negatively influenced hard corals. Colourful *Dendroenphthya* positively influenced *Tubastraea*, which in turn positively influenced encrusting sponges (Fig. 1).

**Figure 1:**
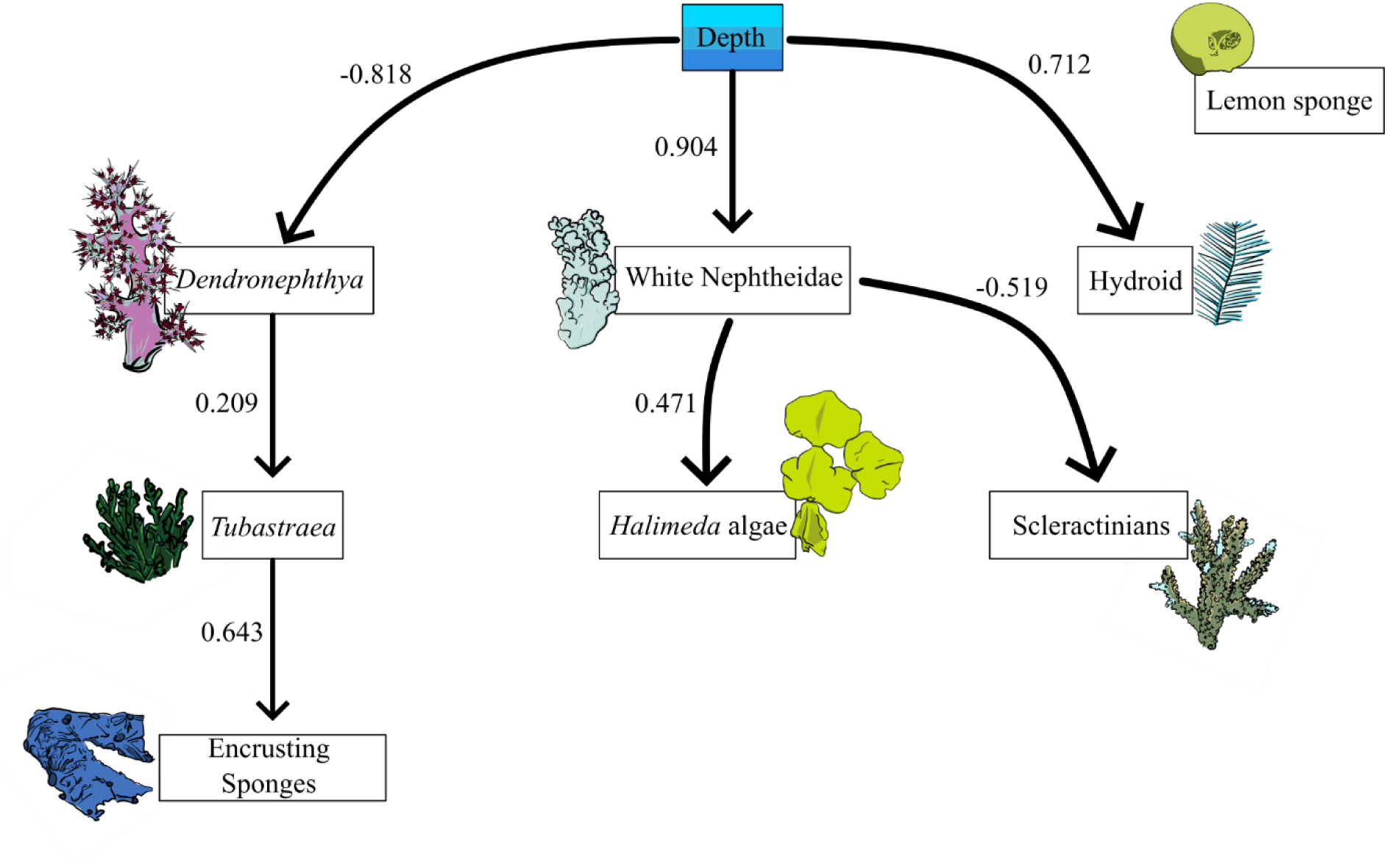
Bayesian network of ecological interactions on the Great White Wall. Thickness of edges reflects occurrence of edges (included in >60% of networks), arrows denote directional effects, numbers next to edges indicate the influence score (strength and sign of interaction).

**Figure 2:**
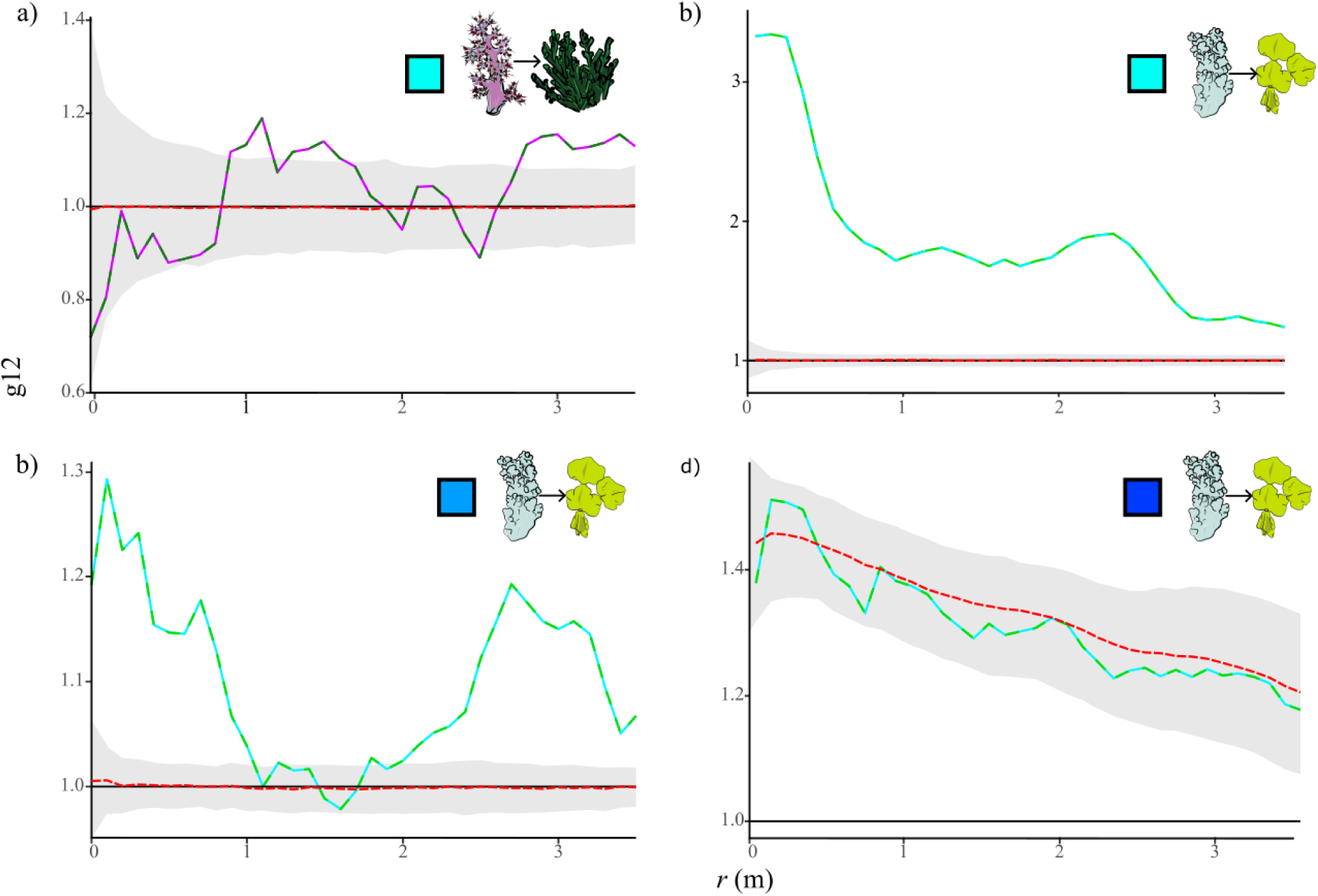
PCF (*y*-axis) plots of bivariate spatial point process analysis for abundant morphospecies identified to directly interact via BNI (Fig. 1) at different depths of the main Great White Wall. a) *Dendronephthya* effect on *Tubastrea* compared to an CSR simulation envelope; b) and c) white Nephtheidae effect on *Halimeda* algae compared to an CSR simulation envelope; d) white Nephtheidae effect on *Halimeda* algae compared to an inhomogeneous linked-Thomas cluster (habitat-mediated mutualism) simulation envelope. Shades of blue indicate depth bin (light is shallow, intermediate is intermediate, dark is deep). Dashed red line indicates fitted model (a-c: CSR; d: ILTC).

### Spatial point process analysis

There were four pairs of taxa that were sufficiently abundant for SPPA: *Dendronephthya* and *Tubastraea* in the shallow mGWW, and white Nephtheidae and *Halimeda* algae at the shallow, intermediate, and deep depth bin of the mGWW. To avoid secondary co-correlations, patial associations were only tested using SPPA in the direction that corresponded to the relationship identified by BNI (*Dendronephthya* on *Tubastraea,* white Nephtheidae on *Halimeda* algae).

We found no best-fit model for the *Dendronephthya-Tubastraea* relationship (Table S3). While the fit for the white Nephtheidae-*Halimeda* algae relationship in the mGWW shallow depth bin was reasonable (*p_d_* = 0.358), visual inspection of the PCF plot indicated that the model fit was actually poor across multiple spatial scales, and so we interpret no model as a good fit for this relationship. We found also no best fit model for the white Nephtheidae-*Halimeda* algae relationship in the mGWW intermediate depth bin. We found a good fit (*p_d_* = 0.474) for an ILTC (habitat-mediated mutualism) model for the white Nephtheidae-*Halimeda* algae relationship for the mGWW deep depth bin indicating a good fit to a mutualistic interaction where the white Nephtheidae facilitate *Halimeda* algae in areas where there is high habitat suitability for *Halimeda* algae.

### Rare morphospecies and diversity analysis

To infer the impact of rarer taxa, our LDA including only rare taxa established three overlapping clusters representing the shallow, intermediate, and deep depth bins across all sites. Note that the BRF was removed due to low sample size after implementation of the testing-training split. The LDA was a good predictor of true depth (64% accuracy). Visual inspection suggested that rare morphospecies diversity was determined by the density of white Nephtheidae (Fig. 3d, red box and arrow).

**Figure 3:**
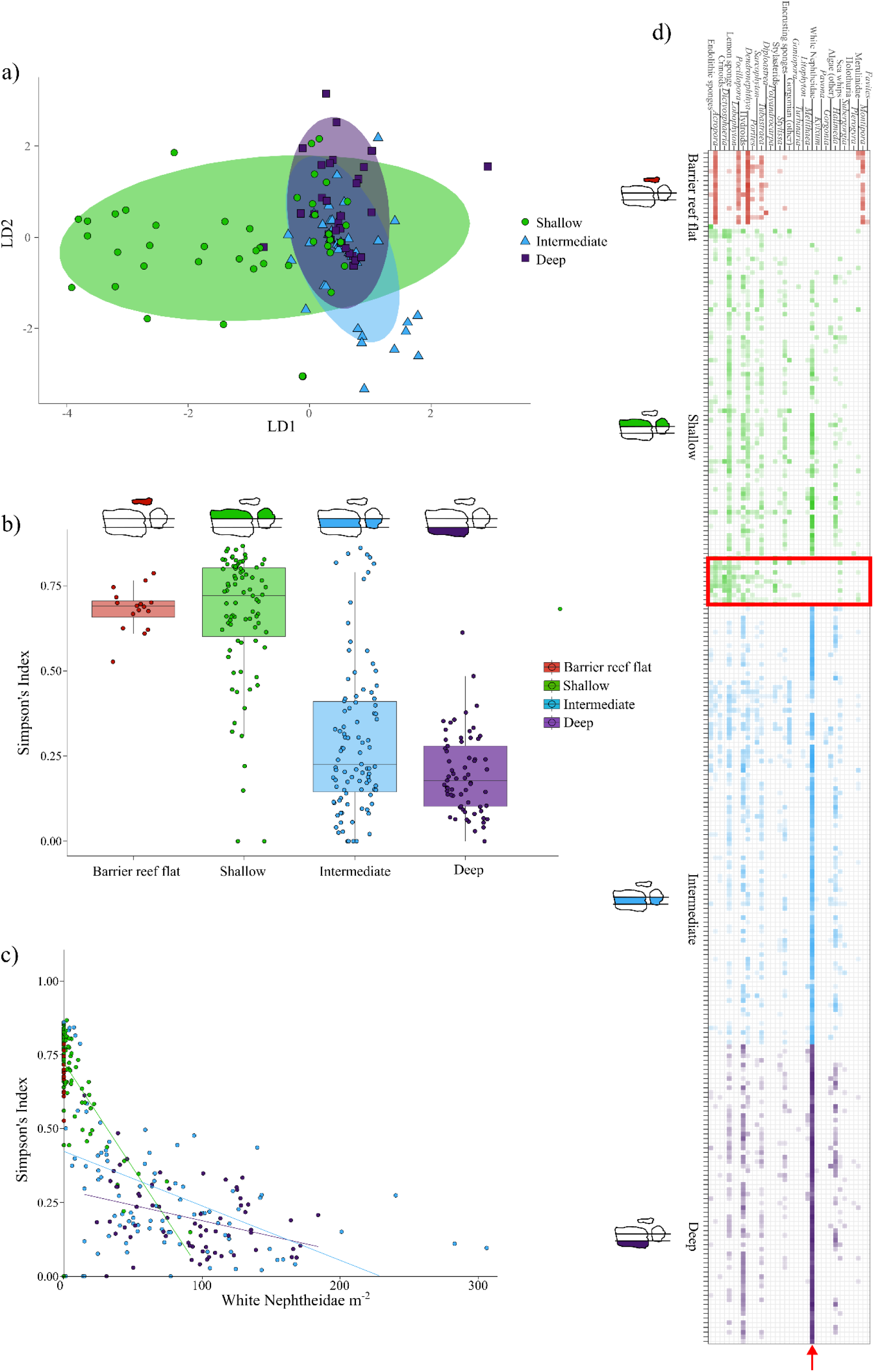
Composition and diversity analysis on the Great White Wall (GWW) and adjacent reefs. On all plots, red = barrier top, green = shallow depth bin, blue = intermediate depth bin, purple = deep depth bin. a) a linear discriminants analysis of the rare taxa, symbols indicate depth bin (circle = shallow, triangle = intermediate, square = deep) and ellipses indicate range occupied by each depth bin; b) differences in Simpson’s diversity index between different depth bins, with points overlayed with a jitter, schematics indicate depth bin; c) association between Simpson’s diversity index with the density of white Nephtheidae, with colours indicating relationships within depth bins; d) matrix plot of morphospecies densities in each quadrat on the GWW and adjacent reefs, the opacity of each cell indicates log density (more opaque = more dense), schematics indicate depth bin. Red box indicates species rich patch with absence of white Nephtheidae soft corals (row indicated by red arrow).

When testing the drivers of diversity for all morphospecies, we found that Simpson, Shannon, and (log transformed) Fisher alpha diversity indices were all negatively associated with increasing depth bin (mean effects on Simpson’s index: Shallow: −0.002; Intermediate: −0.253; Deep: −0.309; Fig. 3b) but 95% confidence intervals crossed zero for all depth bins for all diversity indices apart from the deep depth bin, which had a significant negative effect on Shannon’s index (Table S4). We also found that there was an overall negative association between Simpson, Shannon, and (log transformed) Fisher diversity indices and the density of white Nephtheidae (mean effect Simpson’s index −0.004) and that 95% confidence intervals did not cross zero indicating that the white Nephtheidae may be the main driver of rare taxa diversity (Fig. 3c; Table S4).

## Discussion

In this study, we investigated the ecological interactions between morphospecies across three sites on the GWW, a soft coral-dominated reef in Fiji, using BNI and SPPA. We found a combined effect of depth and soft coral density influenced ecological and diversity dynamics. Depth was the most influential node in the BN, impacting all morphospecies except for lemon sponges, and this pattern is reflected in previous work showing that depth influenced the population ecology of filter feeders and algae on the mGWW (37). In terms of edges between biotic nodes, *Tubastraea* had a positive influence on encrusting sponges in the BN, but sample size precluded any subsequent spatial analysis. It could be that *Tubastraea* acts as suitable habitat for encrusting sponges as they are known to grow over branching scleractinians (65). *Dendronephthya* had a positive influence on the density of *Tubastraea*, but there was no good fit model for this from SPPA (Fig. 2a, Table S3), so the drivers of this relationship are also unclear. The lemon sponges had no strong association with any variables in the network. Previous work found that lemon sponge density was relatively consistent across all mGWW depth bins (37), and there was a minor effect of depth on lemon sponge reproductive clusters compared to *Halimeda* algae, hydroids, and white Nephtheidae, so lemon sponges may have broader biotic and abiotic tolerances than could be tested here. Scleractinians and white Nephtheidae shared the only negative edge between biotic variables in the network, which may reflect segregation arising from competition or distinct habitat preferences, but these effects could not be tested by SPPA since these morphospecies never spatially overlapped in sufficient abundance (n = 100 each). Competition between soft corals and scleractinians has previously been described via allelopathic precluding of settling of scleractinians, which could be a cause of the segregation of the two morphospecies between the GWW sites (66) but large-scale habitat preferences (e.g., variation in light, substrate) cannot be ruled out.

While previous work has highlighted competition between algae and scleractinian corals (67), we found no algae and scleractinian coral edges in our network. The lack of direct association could be a reflection of relatively low impact of human disturbance on the GWW, which has been shown to modulate coral-algae competition (68). We instead found a positive influence of white Nephtheidae soft corals on *Halimeda* algae, and a negative impact of white Nephtheidae on scleractinians, resulting in indirect segregation between scleractinians and algae mediated by soft corals. Subsequent SPPA showed that the best fit model for the white Nephtheidae-algae interaction indicated facilitation and habitat associations (inhomogeneous linked-Thomas cluster process). While it is not possible to disentangle the cause of this mutualism using our data, this unidirectional facilitation could be the algae growing in areas where soft corals have suppressed grazing activity via production of toxic compounds (69,70) or protection mutualisms imparted by invertebrates such as pistol shrimp (71) (Fig. S2). In the intermediate and shallow depth bin, no best-fit model could be found which maybe due to the superposition of changing ecological dynamics (34) with depth or habitat variability (as shown on the mGWW in (37)) or independence because of a lower density of white Nephtheidae. Interestingly, *Halimeda* algae were not found on the AWR where white Nephtheidae were present but in lower densities (although other species of algae were present in low abundances). This difference could reflect a range of ecological interactions such as: i) that the facilitation by the white Nephtheidae is not strong enough on the AWR, ii) a dispersal limitation of the *Halimedia* algae, iii) that the habitat for algae was of a poorer quality, and/or iv) that the relationship is non-monotonic (e.g., thesholded density-dependence (72,73)).

Although the white Nephtheidae soft corals have a positive relationship with algae, there is an overall negative effect of the density of white Nephtheidae on the diversity of any 2m^2^ quadrat in which they are found, indicating that these soft corals may suppress benthic reef diversity. While previous work suggested that soft corals may elevate benthic diversity through temporal niche partitioning whereby intervals of feeding could yield periods where filter feeding benthos can make use of resources in low-current conditions (37), compared to scleractinian reefs, the benthos was significantly less diverse where white Nephtheidae density was high (Fig. 3c). This relationship could, in part, be due to the biodiversity generated by the three-dimensional microhabitat engineered by scleractinians (22). While soft corals also possess large three-dimensional bodies, the structural habitat they generate is ephemeral due to changes in their morphology during periods of low current (25). This transient structural complexity could reduce nursery, sheltering, or wave buffering potential compared to scleractinian-dominated reefs. It is possible that a reduction in grazers or corallivores suppresses the biodiversity benefits generated through intermediate disturbance since soft corals are resilient to removal by wave action, fast currents, or physical removal by reef predators while hunting (as seen with scleractinians and reef sharks (74,75)). Previous work has likewise reported that fish diversity is lower in soft coral-dominated reefs compared to scleractinian reefs (19), although see (76). We found that the composition of rare taxa varies with depth, which could be driven by white Nephtheidae density. We also found that depth was a key contributor to this loss of diversity, which itself is correlated within increasing density of white Nephtheidae, and it is challenging to disentangle the causative element.

Our work highlights the joint role of environment and species interactions in determining reef composition and diversity, in addition to the role of dispersal limitation (37,77). We show that soft coral reefs are composed of a different, and less diverse, assemblage of benthic taxa, than adjacent hard coral reefs, which is concerning given that future climate change may promote soft coral abundance on coral reefs (20). Depth was a key environmental variable in our network, promoting the transition from photosymbiont-hosting scleractinians to aphotosynthetic filter feeder communities. We also show that the mutualistic relationship between white Nephtheidae and algae, which could be due to reducing grazing pressure, occurs most strongly where the white Nephtheidae dominate. Finally, our work suggests that, should soft corals become more prevalent on coral reefs in future climate scenarios, this could lead to a reduction in benthic diversity because soft coral reef systems are underpinned by different ecological dynamics.

## Data accessibility statement

The R code to reproduce this work is available at https://github.com/nis38/GWW_BNI.git and the data are available at https://figshare.com/s/b1de962365bd81568c95.

## Acknowledgements

We thank the Yavusa Mabuco for access to their traditional fishing ground to conduct this research. We thank M. Walser for logistical support.

## Authors’ contributions

NPS: conceptualisation, data curation, formal analysis, methodology, writing draft, revision of manuscript; KMD: data curation, revision of manuscript; CGK: data curation, revision of manuscript; TMK: methodology, revision of manuscript; LK: methodology, revision of manuscript; JW: data curation, revision of manuscript; AAB: data curation, revision of manuscript; VEB: data curation, revision of manuscript; EGM: conceptualisation, data curation, methodology, writing draft, revision of manuscript; AM: conceptualisation, data curation, methodology, writing draft, revision of manuscript.

## Supplementary material

**Figure S1:**
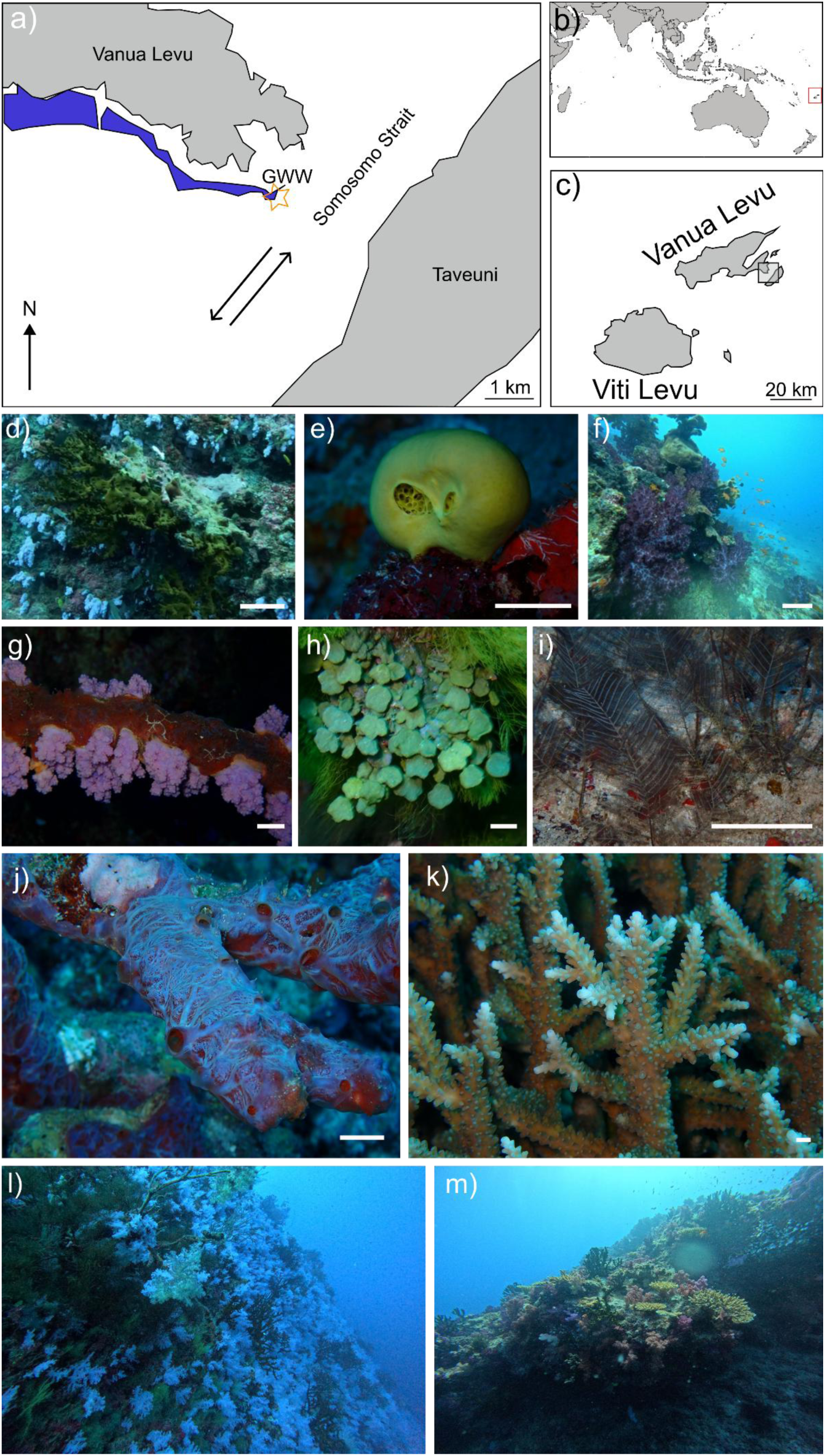
Great White Wall (GWW) and associated reefs localities and morphospecies. a) location of GWW in the Somosomo Strait, Fiji; global location in b), relative to Viti Levu and Vanua Levu islands, Fiji, in c). Arrows represent current direction; blue area represents barrier reef. Abundant reef taxa: d) *Tubastraea* on the GWW, e) Lemon sponge, f) *Dendronephthya* on the barrier reef above the GWW, g) White Nephtheidae on the GWW, h) *Halimeda* algae, i) hydroid colonies, j) encrusting sponge, k) example of scleractinian, likely *Acropora*. Scale bars for e), g), h), i), j), and k) ∼ 1 cm; scale bars of d) and f) ∼ 10 cm. Example reefs: l) the GWW view from ∼ 17 m, and m) the hard and soft coral dominated barrier reef at the top of the GWW at ∼ 12 m.

**Figure S2:**
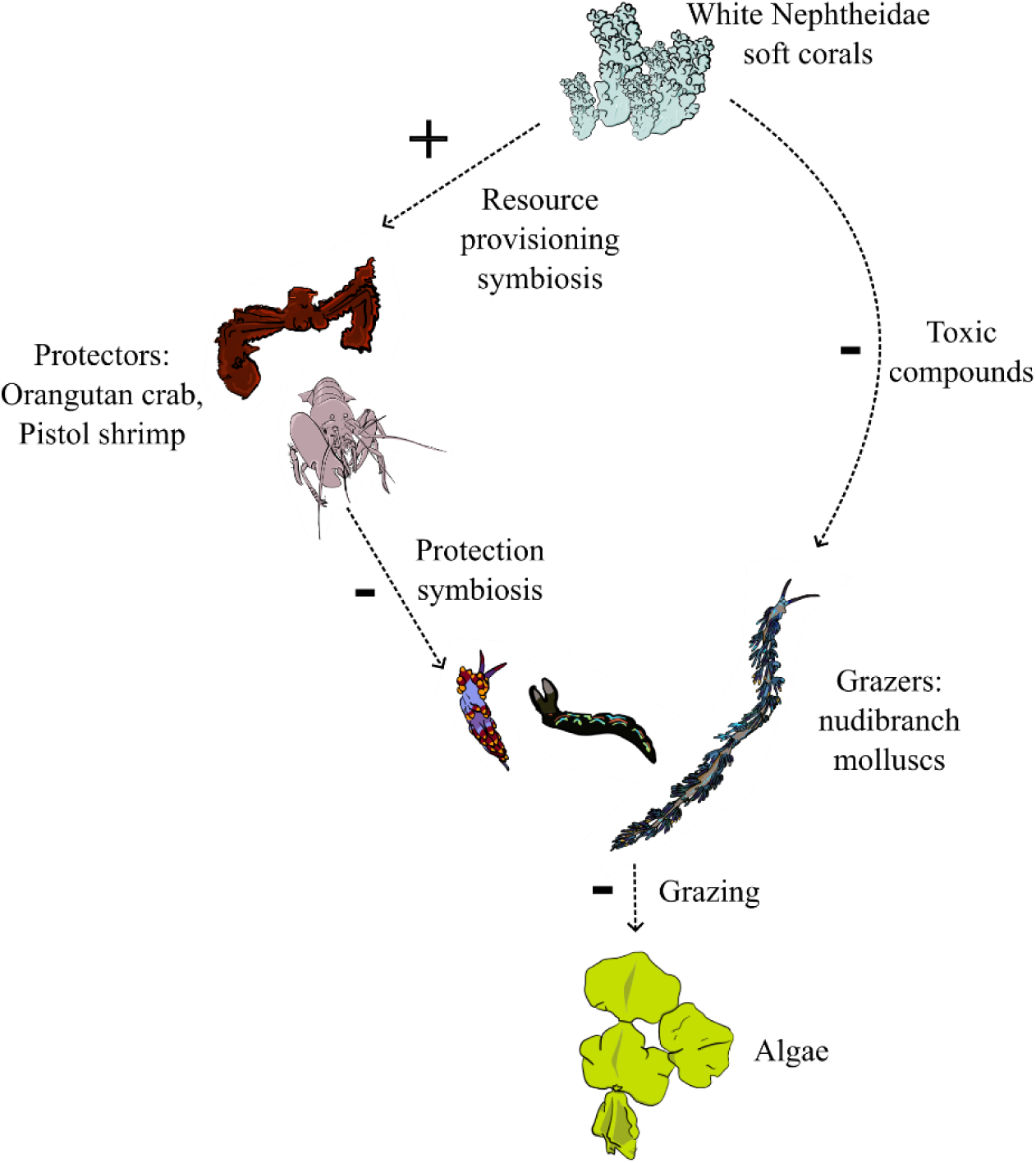
Putative network schematic of plausible mutualism between white Nephtheidae soft corals and *Halimeda* algae (Fig. 1, 2d). White Nephtheidae deploy toxic compounds or facilitate protective mutualists to deter grazers, depressing the local impact of grazing. The release of grazing pressure generates favourable conditions of algal growth, so white Nephtheidae and algae are more commonly found together. We observed all organisms figured here on or near the Great White Wall, apart from the pistol shrimp, which has been found to be associated with Nephtheidae soft corals (71).

**Table S1:** Sample sizes for all morphospecies that appeared frequently enough to be included in Bayesian network analysis. BRF = Barrier reef flat, GWW = Great White Wall, AWR = Adjacent wall reef.

| Morphospecies | BRF | GWW |  |  | AWR |  | Total |
| --- | --- | --- | --- | --- | --- | --- | --- |
|  |  | Shallow | Intermediate | Deep | Shallow | Deep |  |
|  |  |  |  | 10,94 |  |  | 24,539 |
| White Nephthidae | 0 | 1,417 | 9,355 | 3 | 6 | 2,818 |  |
| *Dendronephthya* | 348 | 153 | 86 | 23 | 28 | 55 | 693 |
| *Tubastraea* | 32 | 128 | 94 | 73 | 9 | 16 | 352 |
| Encrusting sponges | 2 | 99 | 62 | 38 | 7 | 6 | 214 |
| Lemon sponges | 2 | 260 | 93 | 96 | 156 | 130 | 737 |
| Halimeda algae | 0 | 155 | 462 | 329 | 0 | 0 | 946 |
| Hydroids | 6 | 159 | 316 | 584 | 55 | 128 | 1,248 |
| Scleractinian | 342 | 22 | 13 | 11 | 98 | 24 | 510 |

**Table S2:** Discretisation splits for all nodes included in Bayesian network analysis. If the %0 value was <40%, the node was discretised into zeros, low abundance, and high abundance, if %0 > 40%, we split groups into only presence and absence, and if %0 > 70%, the node was eliminated from analysis. Note that the environmental variable depth was split into depth bins.

| Node | % 0s | Split |
| --- | --- | --- |
| Depth | 0 | Barrier reef top, shallow wall, intermediate wall, deep wall |
| White Nephthidae | 21 | Zero, low, high |
| Hydroids | 50 | Presence, absence |
| Halimeda Algae | 60 | Presence, absence |
| *Tubastraea* | 45 | Presence, absence |
| *Dendronephthya* | 47 | Presence, absence |
| Scleractinian | 68 | Presence, absence |
| Encrusting sponges | 64 | Presence, absence |
| Lemon sponges | 32 | Zero, low, high |

**Table S3:**
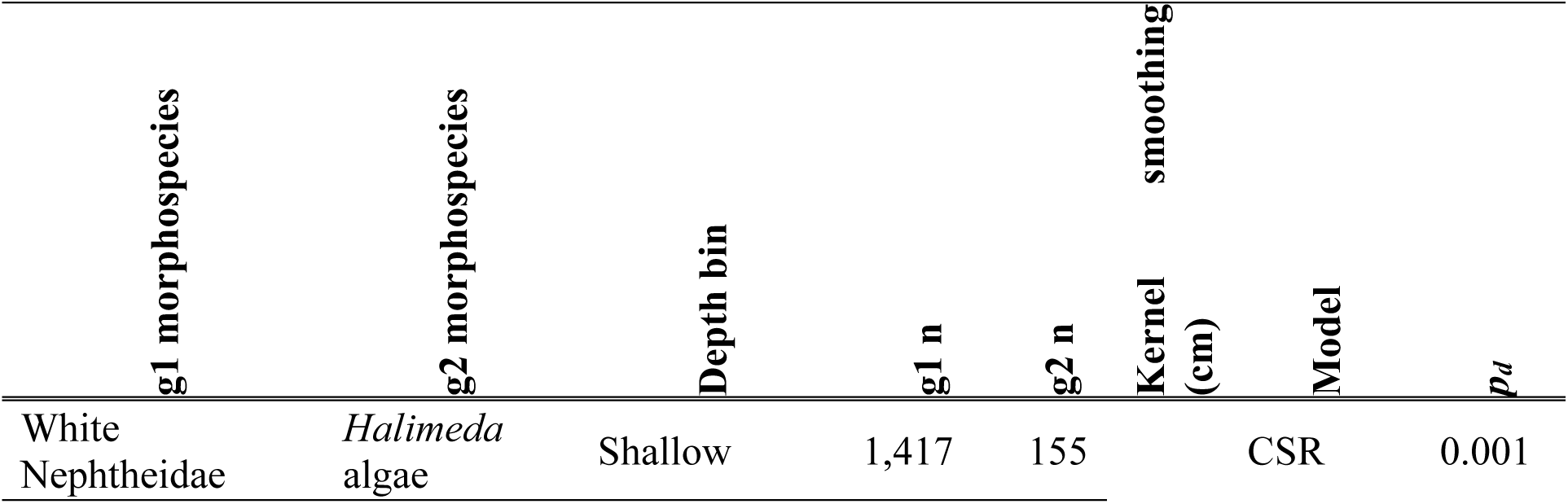
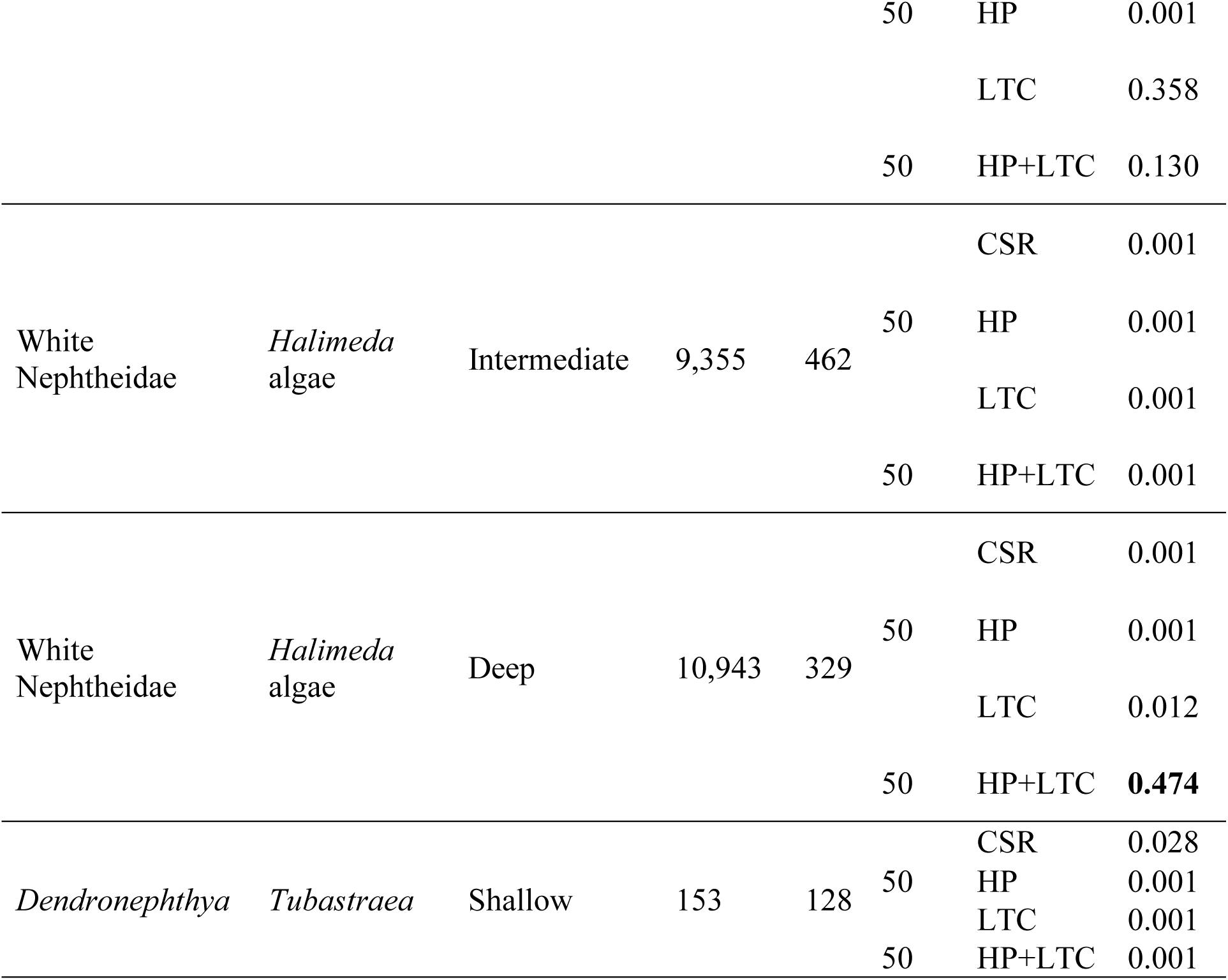
Full model results for all Great White Wall morphospecies that met the requirements for bivariate spatial analysis across best-fit spatial scales test. CSR = complete spatial randomness, HP = heterogeneous Poisson, LTC = Linked Thomas cluster, g1 = identity of pattern 1, g2 = identity of pattern 2. Note that clustering spatial analyses investigate the clustering of g2 around g1, and the HP+LTC models use a HP process generated by the density of g2, whereas HP processes alone use a HP process generated by the density of g1 and g2. All models were fit to a spatial scale from 0 – 350 cm. Bold numbers indicate best-fit model according to Diggle’s goodness of fit (*p_d_*).

**Table S4:**
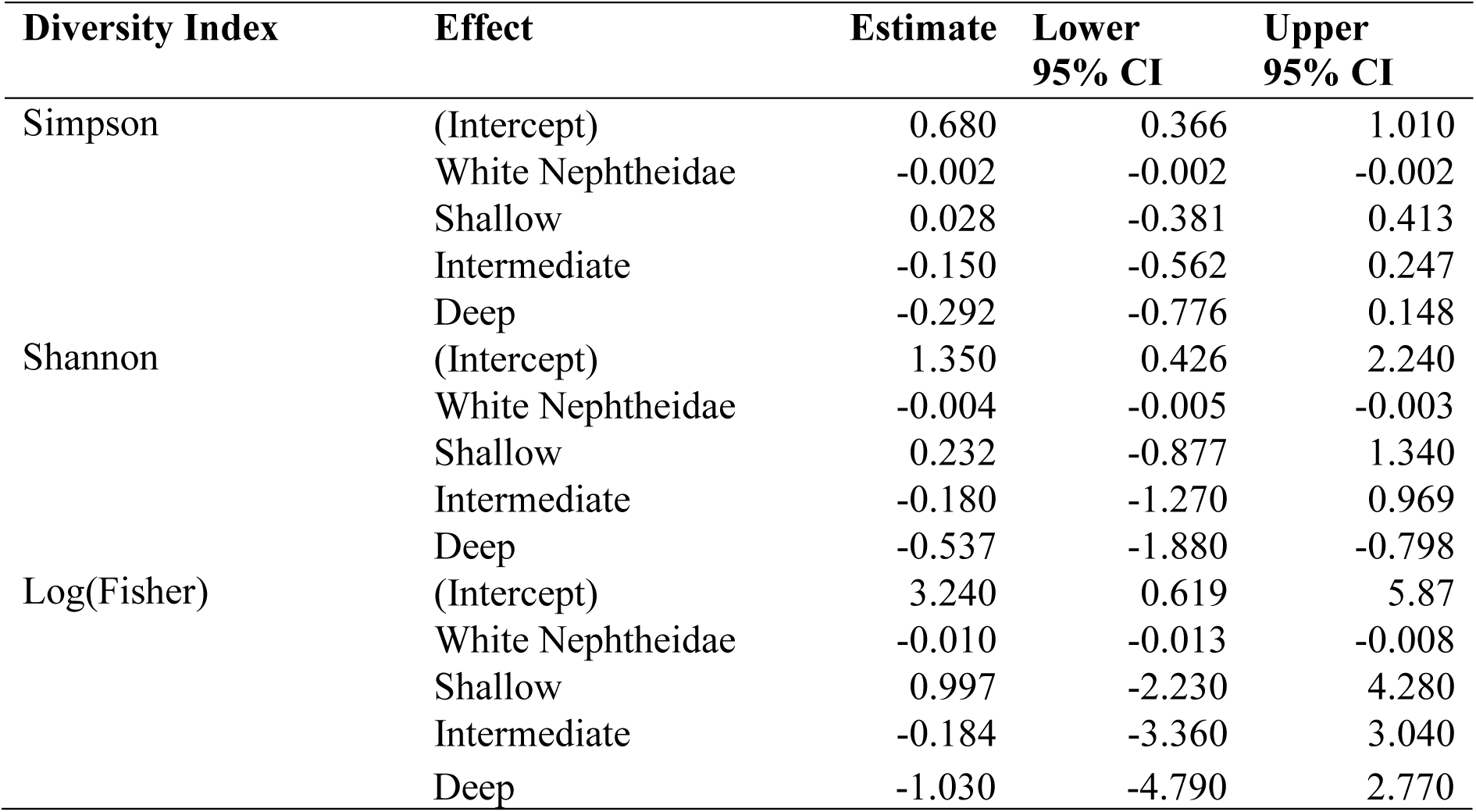
Model coefficients for a linear mixed model of Simpson, Shannon, and (log transformed) Fisher diversity indices and the density of white Nephtheidae and depth, with site as a fixed effect.

## Supplementary methods

The BNI algorithm required each variable to be discretised and used uniform priors to maximise statistical power and mask noise (28,42). Morphospecies were discretised depending on the zero-weighting in the dataset:

1. If >70% of quadrats contained 0s for a given morphospecies, the morphospecies was re-grouped into an ecologically similar morphospecies *cf.* (27–29,43,44). If there was no ecologically similar morphospecies for re-grouping, then the morphospecies were considered too rare for BNI and SPPA.
2. If <70% and >40% of quadrats contained 0s for a given morphospecies and there was no other sensible re-grouping, the morphospecies was assigned presence-absence.
3. If <40% of quadrats contained 0s for a given morphospecies, the morphospecies was assigned either a zero, low, or high count, with low and high counts split by the median density across all quadrats after removing 0s.

BNs cannot normally be analytically found so a search algorithm was required to find the best-fit BN which used BIC to find the best performing network *cf.* (28,43). A greedy search (maximising nodes) with hill climbing was implemented *cf.*(29). To minimise outlier bias, each search used a data set of 1,000 samples with a 95% bootstrap (with replacement) *cf.* (28). Edge frequencies were bimodally distributed and to define high and low probability edges. For each edge, the directionality was the direction which occurred more often in BNs (>50% ± 10%). We averaged the best 10 networks and only the high frequency edges (>0.6) contributed to the network, with their influence score (IS) calculated as a mean of the constituent edge IS (30).

For the spatial analysis, since the only edges that met the sample size requirements were positive, we tested the aggregation models only, namely : i) homogenous Poisson, or complete spatial randomness (CSR), indicating no interactions over the spatial scales tested (34,49); ii) a heterogeneous Poisson (HP) process, modelling aggregation due to habitat associations; iii) a linked-Thomas cluster (LTC) process, modelling aggregation due to mutualisms (29,50–52); and iv) an inhomogenous linked-Thomas cluster (ILTC) process, modelling mutualisms in areas of high habitat suitability. For HP analysis, habitat heterogeneity can be approximated by smoothing the density map of a morphospecies (53). We produced density maps of each morphospecies using an Epanechnikov kernel with a bandwidth of 0.5 m, a value that consistently provided a balance between the locations of the organisms and the variation generated by them *cf.* (29). To test whether a bivariate PCF was best fit to a given null model PCF, 999 Monte Carlo simulations were run for CSR, HP, LTC, and ILTC processes, and the simulation envelopes taken to be between the 5% highest and lowest values *cf.* (34). CSR and HP models were fit using maximum likelihood methods and LTC and ILTC models were fit using minimum contrast methods (54,55).

A goodness-of-fit test was then used to quantitatively assess differences between the observed pattern PCFs and simulated PCFs. Goodness-of-fit tests provide a hypothesis test, with the null hypothesis being that the measured process (here, PCF for each pair of morphospecies’ point patterns) departs from the simulation envelope over a specified distance interval (34,56). Here, we used Diggle’s goodness-of-fit test (*p_d_*), with high *p_d_* (i.e., closer to 1) interpreted to be a good model fit (56), alongside visual inspection of the morphospecies’ PCF plot (34).

## References

1. Roberts CM, McClean CJ, Veron JEN, Hawkins JP, Allen GR, McAllister DE, et al. Marine Biodiversity Hotspots and Conservation Priorities for Tropical Reefs. Science. 2002 Feb 15;295(5558):1280–4.

2. Graham NAJ, Bellwood DR, Cinner JE, Hughes TP, Norström AV, Nyström M. Managing resilience to reverse phase shifts in coral reefs. Front Ecol Environ. 2013 Dec;11(10):541–8.

3. Hughes TP, Barnes ML, Bellwood DR, Cinner JE, Cumming GS, Jackson JBC, et al. Coral reefs in the Anthropocene. Nature. 2017 Jun;546(7656):82–90.

4. Hughes TP, Anderson KD, Connolly SR, Heron SF, Kerry JT, Lough JM, et al. Spatial and temporal patterns of mass bleaching of corals in the Anthropocene. Science. 2018 Jan 5;359(6371):80–3.

5. Reimer JD, Peixoto RS, Davies SW, Traylor-Knowles N, Short ML, Cabral-Tena RA, et al. The Fourth Global Coral Bleaching Event: Where do we go from here? Coral Reefs. 2024 Aug;43(4):1121–5.

6. Inoue S, Kayanne H, Yamamoto S, Kurihara H. Spatial community shift from hard to soft corals in acidified water. Nat Clim Change. 2013 Jul;3(7):683–7.

7. Toledo-Rodriguez DA, Veglia AJ, Marrero NMJ, Gomez-Samot JM, McFadden CS, Weil E, et al. Shadows over Caribbean reefs: occurrence of a new invasive soft coral species, Xenia umbellata, in southwest Puerto Rico. Coral Reefs [Internet]. 2025 May 12 [cited 2025 Jun 16]; Available from: https://link.springer.com/10.1007/s00338-025-02670-5

8. Anton A, Randle JL, Garcia FC, Rossbach S, Ellis JI, Weinzierl M, et al. Differential thermal tolerance between algae and corals may trigger the proliferation of algae in coral reefs. Glob Change Biol. 2020 Aug;26(8):4316–27.

9. Carballo-Bolaños R, Soto D, Chen CA. Thermal Stress and Resilience of Corals in a Climate-Changing World. J Mar Sci Eng. 2019 Dec 24;8(1):15.

10. Krieger EC, Taise A, Nelson WA, Grand J, Le Ru E, Davy SK, et al. Tolerance of coralline algae to ocean warming and marine heatwaves. Li M, editor. PLOS Clim. 2023 Jan 4;2(1):e0000092.

11. Lopes AR, Faleiro F, Rosa IC, Pimentel MS, Trubenbach K, Repolho T, et al. Physiological resilience of a temperate soft coral to ocean warming and acidification. Cell Stress Chaperones. 2018 Sep;23(5):1093–100.

12. Fabricius KE. Soft coral abundance on the central Great Barrier Reef: effects of Acanthaster planci, space availability, and aspects of the physical environment. Coral Reefs. 1997 Jul;16(3):159–67.

13. Lalas JAA, Manzano GG, Desabelle LAB, Baria-Rodriguez MV. Spatial variation in the benthic community structure of a coral reef system in the central Philippines: Highlighting hard coral, octocoral, and sponge assemblages. Reg Stud Mar Sci. 2023 Jul;61:102859.

14. Bastidas C, Fabricius KE, Willis BL. Demographic aspects of the soft coral Sinularia flexibilis leading to local dominance on coral reefs. 2004;

15. Joshi JD, Joshi DM, Patel RS, Munjpara SB, Salvi HD. A record of Mono-specific Carpets of Genus –Sinulariaon Coral reefs of the Gulf of Kachchh, Gujarat, India. Res J Biol Sci. 2015;4.

16. Larkin MF, Davis TR, Harasti D, Cadiou G, Poulos DE, Smith SDA. The rapid decline of an Endangered temperate soft coral species. Estuar Coast Shelf Sci. 2021 Jul;255:107364.

17. Coffroth MA, Buccella LA, Eaton KM, Lasker HR, Gooding AT, Franklin H. What makes a winner? Symbiont and host dynamics determine Caribbean octocoral resilience to bleaching. Sci Adv. 2023 Nov 24;9(47):eadj6788.

18. Thobor B, Tilstra A, Bourne DG, Springer K, Mezger SD, Struck U, et al. The pulsating soft coral Xenia umbellata shows high resistance to warming when nitrate concentrations are low. Sci Rep. 2022 Oct 6;12(1):16788.

19. Lalas JAA, Gomez R, Abram A, Hakim AA, Nakamura T, Reimer JD. Patterns of fish assemblage structure on reefs with varying degrees of hard coral and soft coral dominance in Okinawa Island, Japan. Mar Biodivers. 2024 Dec;54(6):82.

20. Tsounis G, Edmunds PJ. Three decades of coral reef community dynamics in St. John, USVI : a contrast of scleractinians and octocorals. Ecosphere. 2017 Jan;8(1):e01646.

21. Eddy TD, Lam VWY, Reygondeau G, Cisneros-Montemayor AM, Greer K, Palomares MLD, et al. Global decline in capacity of coral reefs to provide ecosystem services. One Earth. 2021 Sep;4(9):1278–85.

22. Woodhead AJ, Hicks CC, Norström AV, Williams GJ, Graham NAJ. Coral reef ecosystem services in the Anthropocene. Fox C, editor. Funct Ecol. 2019 Jun;33(6):1023–34.

23. Sandin SA, McNamara DE. Spatial dynamics of benthic competition on coral reefs. Oecologia. 2012 Apr;168(4):1079–90.

24. Romero GQ, Gonçalves-Souza T, Vieira C, Koricheva J. Ecosystem engineering effects on species diversity across ecosystems: a meta-analysis. Biol Rev. 2015 Aug;90(3):877– 90.

25. Hellström M, Benzie JAH. Robustness of size measurement in soft corals. Coral Reefs. 2011 Sep;30(3):787.

26. Hammond J, Smith VA. Bayesian networks for network inference in biology. J R Soc Interface. 2025;22.

27. Khan TM, Griffiths HJ, Whittle RJ, Stephenson NP, Delahooke KM, Purser A, et al. Network analyses on photographic surveys reveal that invertebrate predators do not structure epibenthos in the deep (∼2000m) rocky Powell Basin, Weddell Sea, Antarctica. Front Mar Sci. 2024 Jul 2;11:1408828.

28. Milns I, Beale CM, Smith VA. Revealing ecological networks using Bayesian network inference algorithms. Ecology. 2010 Jul;91(7):1892–9.

29. Mitchell EG, Harris S. Mortality, Population and Community Dynamics of the Glass Sponge Dominated Community “The Forest of the Weird” From the Ridge Seamount, Johnston Atoll, Pacific Ocean. Front Mar Sci. 2020 Oct 20;7:565171.

30. Yu J, Smith VA, Wang PP, Hartemink AJ, Jarvis ED. Advances to Bayesian network inference for generating causal networks from observational biological data. Bioinformatics. 2004 Dec 12;20(18):3594–603.

31. Mitchell EG, Butterfield NJ. Spatial analyses of Ediacaran communities at Mistaken Point. Paleobiology. 2018 Feb;44(1):40–57.

32. McFadden IR, Bartlett MK, Wiegand T, Turner BL, Sack L, Valencia R, et al. Disentangling the functional trait correlates of spatial aggregation in tropical forest trees. Ecology. 2019 Mar;100(3):e02591.

33. Velázquez E, Martínez I, Getzin S, Moloney KA, Wiegand T. An evaluation of the state of spatial point pattern analysis in ecology. Ecography. 2016 Nov;39(11):1042–55.

34. Wiegand T, Moloney KA. Handbook of Spatial Point-Pattern Analysis in Ecology [Internet]. 1st Edition. CRC; 2013. Available from: 10.1201/b16195

35. Raventós J, Wiegand T, Luis MD. Evidence for the spatial segregation hypothesis: a test with nine-year survivorship data in a Mediterranean shrubland. Ecology. 2010 Jul;91(7):2110–20.

36. Illian JB, Penttinen A, Stoyan H, Stoyan D. Statistical analysis and modelling of spatial point patterns. John Wiley & Sons; 2008.

37. Stephenson NP, Delahooke KM, Kenchington CG, Waitaiti J, Ball AA, Bonito VE, et al. Depth affects the population dynamics on a soft coral-dominated reef on the Great White Wall, Fiji. Coral Reefs. 2025;

38. Tomar S. Converting video formats with FFmpeg. Belltown Media; 2006.

39. Delahooke KM, Liu AG, Stephenson NP, Mitchell EG. ‘Conga lines’ of Ediacaran fronds: insights into the reproductive biology of early metazoans. R Soc Open Sci. 2024 May;11(5):231601.

40. R Core Team. R: A language end environment for statistical computing [Internet]. Vienna, Austria: R Foundation for Statistical Computing; 2025. Available from: https://www.R-project.org/

41. Smith VA, Yu J, Smulders T, Hartemink AJ, Jarvis ED. Computational Inference of Neural Information Flow Networks. PLoS Comput Biol. 2006;preprint(11):e161.

42. Yu J, Smith VA, Wang PP, Hartemink AJ, Jarvis ED. Using Bayesian network inference algorithms to recover molecular genetic regulatory networks. Int Conf Syst Biol. 2002;2002.

43. Mitchell EG, Durden JM, Ruhl HA. First network analysis of interspecific associations of abyssal benthic megafauna reveals potential vulnerability of abyssal hill community. Prog Oceanogr. 2020 Aug;187:102401.

44. Stafford R, Smith VA, Husmeier D, Grima T, Guinn B ann. Predicting ecological regime shift under climate change: New modelling techniques and potential of molecular-based approaches. Curr Zool. 2013 Jun 1;59(3):403–17.

45. Brown C, Law R, Illian JB, Burslem DFRP. Linking ecological processes with spatial and non-spatial patterns in plant communities. J Ecol. 2011 Nov;99(6):1402–14.

46. May F, Huth A, Wiegand T. Moving beyond abundance distributions: neutral theory and spatial patterns in a tropical forest. Proc R Soc B Biol Sci. 2015 Mar 7;282(1802):20141657.

47. Mitchell EG, Kenchington CG, Liu AG, Matthews JJ, Butterfield NJ. Reconstructing the reproductive mode of an Ediacaran macro-organism. Nature. 2015 Aug;524(7565):343–6.

48. Wiegand T. Programita [Internet]. 2018. Available from: https://programita.org/

49. Diggle PJ, Eglen SJ, Troy JB. Modelling the Bivariate Spatial Distribution of Amacrine Cells. In: Baddeley A, Gregori P, Mateu J, Stoica R, Stoyan D, editors. Case Studies in Spatial Point Process Modeling [Internet]. New York: Springer-Verlag; 2006 [cited 2025 Jun 16]. p. 215–33. (Lecture Notes in Statistics; vol. 185). Available from: http://link.springer.com/10.1007/0-387-31144-0_12

50. Dhungana A, Mitchell EG. Facilitating corals in an early Silurian deep-water assemblage. Cherns L, editor. Palaeontology. 2021 May;64(3):359–70.

51. Dickie IA, Schnitzer SA, Reich PB, Hobbie SE. Spatially disjunct effects of co-occurring competition and facilitation. Ecol Lett. 2005 Nov;8(11):1191–200.

52. Lingua E, Cherubini P, Motta R, Nola P. Spatial structure along an altitudinal gradient in the Italian central Alps suggests competition and facilitation among coniferous species. J Veg Sci. 2008 Jun;19(3):425–36.

53. Shen G, Yu M, Hu XS, Mi X, Ren H, Sun IF, et al. Species–area relationships explained by the joint effects of dispersal limitation and habitat heterogeneity. Ecology. 2009 Nov;90(11):3033–41.

54. Baddeley A, Turner R. Practical Maximum Pseudolikelihood for Spatial Point Patterns: (with Discussion). Aust N Z J Stat. 2000 Sep;42(3):283–322.

55. Diggle PJ, Gratton RJ. Monte Carlo Methods of Inference for Implicit Statistical Models. J R Stat Soc Ser B Stat Methodol. 1984 Jan 1;46(2):193–212.

56. Diggle PJ. Statistical Analysis of Spatial and Spatio-Temporal Point Patterns. 1st ed. CRC Press; 2003.

57. Kondratyeva A, Grandcolas P, Pavoine S. Reconciling the concepts and measures of diversity, rarity and originality in ecology and evolution. Biol Rev. 2019 Aug;94(4):1317–37.

58. McGill BJ, Etienne RS, Gray JS, Alonso D, Anderson MJ, Benecha HK, et al. Species abundance distributions: moving beyond single prediction theories to integration within an ecological framework. Ecol Lett. 2007 Oct;10(10):995–1015.

59. Manica A, Carter RW. Morphological and fluorescence analysis of the Montastraea annularis species complex in Florida. Mar Biol. 2000;137:899–906.

60. Simpson EH. Measurement of Diversity. Nature. 1949;163(688).

61. Clarke KR, Warwick RM. Change in Marine COmmunities: An Approach to Statistical Analysis and Interpretation. 2nd Edition. PR IM ER -E; 2001.

62. Shannon CE, Weaver W. The Mathematical Theory of Communication. Urbana: University of Illinois Press; 1949.

63. Fisher RA, Corbet AS, Williams CB. The Relation Between the Number of Species and the Number of Individuals in a Random Sample of an Animal Population. J Anim Ecol. 1943 May;12(1):42.

64. Bates D, Mächler M, Bolker B, Walker S. Fitting Linear Mixed-Effects Models Using **lme4**. J Stat Softw [Internet]. 2015 [cited 2025 Jun 19];67(1). Available from: http://www.jstatsoft.org/v67/i01/

65. Chaves-Fonnegra A, Zea S. Coral colonization by the encrusting excavating Caribbean sponge Cliona delitrix: Cliona delitrix coral colonization. Mar Ecol. 2011 Jun;32(2):162– 73.

66. Maida M, Sammarco PW, Coil JC. Effects of soft corals on scleractinian coral recruitment. I: Directional allelopathy and inhibition of settlement. Mar Ecol Prog Ser. 1995;121:191–202.

67. McCook L, Jompa J, Diaz-Pulido G. Competition between corals and algae on coral reefs: a review of evidence and mechanisms. Coral Reefs. 2001 May;19(4):400–17.

68. Barott K, Williams G, Vermeij M, Harris J, Smith J, Rohwer F, et al. Natural history of coral−algae competition across a gradient of human activity in the Line Islands. Mar Ecol Prog Ser. 2012 Jul 24;460:1–12.

69. Ayllon ME, Elpa HES, Metillo EB, Uy MM. Toxicity of crude extracts from soft corals (Anthozoa, Alcyonacea) collected at varying wave exposure sites in Talisayan, Northern Mindanao, Philippines. 2019;

70. Kasumyan AO, Tinkova TV. Spreading of deterrency as a means of chemical defense among aquatic organisms inhabiting the coral reefs of Vietnam. Dokl Biol Sci. 2014 Jan;454(1):39–42.

71. Nakano R, Fujii T. The soft-coral associated pistol shrimp Synalpheus neomeris (De Man) (Decapoda: Alpheidae) defends its host against nudibranchs in Okinawa, Japan. RAFFLES Bull Zool. 2014;

72. Aizen MA, Morales CL, Vázquez DP, Garibaldi LA, Sáez A, Harder LD. When mutualism goes bad: density-dependent impacts of introduced bees on plant reproduction. New Phytol. 2014 Oct;204(2):322–8.

73. Zhang Z. Mutualism or cooperation among competitors promotes coexistence and competitive ability. Ecol Model. 2003 Jun;164(2–3):271–82.

74. Jiménez-Centeno CE. Coral colony fragmentation by whitetip reef sharks at Coiba Island National Park, Panama. Rev Biol Trop. 1997;45(1):698–700.

75. Roff G, Doropoulos C, Rogers A, Bozec YM, Krueck NC, Aurellado E, et al. The Ecological Role of Sharks on Coral Reefs. Trends Ecol Evol. 2016 May;31(5):395–407.

76. Epstein HE, Kingsford MJ. Are soft coral habitats unfavourable? A closer look at the association between reef fishes and their habitat. Environ Biol Fishes. 2019 Mar;102(3):479–97.

77. Quattrini AM, Gómez CE, Cordes EE. Environmental filtering and neutral processes shape octocoral community assembly in the deep sea. Oecologia. 2017 Jan;183(1):221– 36.

